# Immobilized Nucleoside 2′-Deoxyribosyltransferases from Extremophiles for Nucleoside Biocatalysis

**DOI:** 10.1101/2024.09.20.614042

**Authors:** Saúl Antonio Hernández Martínez, Peijun Tang, Roberto Parra-Saldívar, Elda M. Melchor-Martínez, Clarissa Melo Czekster

## Abstract

The synthesis of nucleosides is crucial for pharmaceutical and biotechnological applications, acting as drugs and as essential building blocks for numerous therapeutic agents. However, most enzymes employed in nucleoside biocatalysis are not recycled, possess limit stability and strict substrate selection for ribonucleosides or 2′deoxyribonucleosides. We employed 2′deoxyribonucleoside transferase (NDT) enzymes from thermophilic and psychrophilic bacteria to demonstrate they can be immobilized to en-hance specific activity, stability, and recyclability. NDT enzymes from *Chroococcidiopsis thermalis* (CtNDT), and *Bacillus psychrosaccharolyticus* (BpNDT) were immobilized by covalent attachment to chitosan beads. A double mutant of CtNDT, capable of generating 3′deoxyribonucleosides, showed remarkable and increased stability after immobilization when compared to the same enzyme in solution. Furthermore, we demonstrated the recyclability of immobilized biocatalysts, with a tenfold improvement in reaction yield over 20 consecutive cycles, highlighting the practicality and sustainability of the developed immobilization method. We used our strategy to produce a pharmaceutically relevant 3′deoxyribonucleoside (2-fluoro-3′-deoxyadenosine). This highlights the importance of efficient immobilization techniques to enhance catalytic properties of NDT enzymes, expanding their utility in biocatalysis.

## 1. INTRODUCTION

Nucleosides are the building blocks of all nucleic acids, playing pivotal roles in various cellular processes such as enzyme regulation, metabolism, DNA/RNA synthesis, and signaling cascades. Therefore, nucleoside analogues (NAs) possess activity against viruses, cancer, bacteria, and as starting materials for antisense oligonucleotides^1^. Currently more than 90 nucleoside and nucleotide-based drugs are approved for the treatment of different infections, and cancers (e.g. idoxuridine, brivudine, cytarabine).^2^ The success of NAs as drugs can be attributed to extensive efforts to isolate novel nucleoside natural products and to advances in organic synthesis methods.^3^ However, both present limitations, as natural product isolation can be complex and low-yielding^4^, while synthesis frequently utilizes harsh conditions, requires selective protection of polar hydroxy and amino groups and is further complicated by limited solubility of nucleosides in most organic solvents.^5^ Moreover, both might also require time-consuming multistep processes, increasing production of undesired byproducts and decreasing the overall synthesis efficiency.^1,6^ Biocatalysis presents an attractive alternative, being more sustainable^7^, eliminating protection/deprotection steps and relying on the outstanding catalytic power of enzymes to generate fewer (or no) side products^8^.

Biocatalytic methods for chemo-enzymatic synthesis of organic molecules have been amply discussed elsewhere.^9–11^ These methods include whole cell biocatalysis using bacteria or fungi^12^ or isolated enzymes^13^, both resulting in substantial conversion rates. However, a drawback of using whole-cell or purified enzymes lies in their low or nonexistent recyclability, significantly increasing production costs.^14^ Enzyme immobilization is a strategy to overcome these limitations since strategies employing covalent bonding, entrapment, or encapsulation can significantly enhance the long-term stability of enzymes.^15^ This enhancement provides resistance to degradation or denaturation^16^, and greatly facilitates the separation of biocatalysts from reaction mixtures, including final products, streamlining the purification process.^17^

Nucleoside phosphorylases (NPs) have been extensively employed in biocatalysis as they catalyze the synthesis of nucleosides in a two-step reaction.^18^ However, issues with phosphor-sugar stability and reaction equilibrium present limitations to the production of some NAs, and their engineering is an active area of study to overcome these obstacles.^19^ NPs failed to recognize cytosine derivatives as substrates, requiring a two-enzyme one-pot approach for the production of purine nucleosides.^20^ Nucleoside 2′-deoxyribosyltransferases (dNDTs) are enzymes with strict specificity for 2′- deoxyribonucleosides which have been employed as biocatalysts for synthesizing numerous 2′-deoxyribonucleoside analogues with potential biomedical applications.^21,22^ dNDTs participate in deoxyribonucleoside salvage in some organisms, and catalyze the reaction via a ping-pong mechanism with the formation of a 2’-deoxyribosyl-enzyme intermediate covalently linked to an active-site glutamate residue.^23,24^

Immobilization of NPs^25–30^ and dNDTs^21,31,32^ for biocatalysis faces stability and activity losses as bottlenecks, ultimately affecting reaction overall yield and recyclability. To overcome these issues, we employed the enzyme from the thermophilic *organism Chroococcidiopsis thermalis* PCC 7203 (CtNDT) as well as a double mutant (CtNDT_Y7F_A9S_), and the enzyme from the psychrotolerant organism *Bacillus psychrosaccharolyticus* (*Bp*NDT) in different immobilization techniques including covalent bonding, entrapment, and encapsulation. Immobilization allowed enzyme recycling, increasing nucleoside production, and improving long term operability. Moreover, CtNDT_Y7F_A9S_ can be used in single enzymatic step to produce 3′-deoxyribonucleoside (2-fluoro-3′-deoxyadenosine) with potential use as an anti-trypanossomal compound^33^. We used immobilized CtNDT_Y7F_A9S_ in seven consecutive cycles to produce the desired nucleoside product, significant increasing reaction yield. In summary, this work sets the stage to using immobilized *Ct*NDT variants in the production of novel nucleosides with potential therapeutic applications.

## 2. MATERIALS AND METHODS

### 2.1. Materials

The codon optimized synthetic gene to express the 2′-deoxyribosyltransferases (NDT) in an *Escherichia coli* host was ordered from Integrated DNA Technologies (IDT). General chemicals and reagents were from Fluorochem, Merck and Fisher Scientific. All chemicals used in the work were of the highest available purity and analytical grade, solvents for HPLC and LC-MS were of HPLC grade.

### 2.2. Cloning and expression of the 2′ -deoxyribosyltransferases

The cloning and expression of *Chroococcidiopsis thermalis* NDT (*Ct*NDT), CtNDT double mutant (CtNDT_Y7F_A9S_), and the *Bacillus psychrosaccharolyticus* NDT (BpNDT) were performed as previously reported ^20,23^. The synthetic gene encoding nucleoside 2’-deoxyribosyltransferase from *Chroococcidiopsis thermalis* (Uniprot code: K9TVX3) and *Bacillus psychrosaccharolyticus* NDT (Uniports code: A0A3G5BRZ6) were cloned into a pJ411 (CtNDT, CtNDT_Y7F_A9S_) or pJ414 (BpNDT) expression plasmid with a cleavable 6-histidine tag at N-terminus. Briefly, the NDT species were transformed into *E. coli* BL21 (DE3) cells for overexpression in LB medium with 50 mg/ml kanamycin at 37°C 180 rpm shaking until the cells reached OD_600_ = 0.8. Protein expression was induced by addition of 0.5 mM of IPTG, followed by incubation overnight at 16°C while shaking at 180 rpm.

### 2.3. Enzyme purification

Following growth, cells were harvested by centrifugation at 12,000g for 20 mins, resuspended in wash buffer (50 mM MES, 250 mM NaCl, 30 mM Imidazole, pH 6.5) and lysed using a cell homogenizer (Constant Systems). Cell debris were removed by centrifugation for 30 mins at 51,000g at 4°C, the supernatant was filtered with a 0.8 mm filter to remove particulates and loaded onto a 5 mL HisTrap column pre-equilibrated with wash buffer. The column was washed 10 column volumes wash buffer, and the NDT proteins were eluted with 50 mM MES, 250 mM NaCl, 500 mM Imidazole, pH 6.5. Fractions containing NDT were pooled and dialyzed with 2 mg/ml Tobacco Etch Virus (TEV) protease (produced in house) in 50 mM MES, 250 mM NaCl, pH 6.5 overnight at 4°C. The dialyzed mixture was loaded onto a 5 mL HisTrap column, and the flow-through fractions were collected and analyzed by 15% SDS-PAGE. Fractions containing pure protein (>95%) were pooled, flash frozen and used in the subsequent experiments.^23^ Protein concentrations were measured according to Bradford method using crystalline bovine serum albumin (BSA) as a standard ^34^.

### 2.4. Enzyme immobilization methods

#### 2.4.1. Encapsulation of CtNTD on Alginate Beads

*Ct*NDT was immobilized into Alginate Beads (AlgB) by direct encapsulation applying a gelation method. An alginate solution (3 %) was prepared by dissolving 0.6 g of sodium alginate salt into 20 mL of deionized water and mixed using a magnetic stirrer until a homogeneous solution was obtained. *Ct*NDT immobilization was achieved by mixing 50 μg of the enzyme with 1 mL of the alginate solution under rotational mixing for 30 min. The resulting solution was extruded through a syringe needle (0.8 x 40 mm) into 5 mL of gelation medium (0.3 CaCl_2_) under slow magnetic stirring to avoid the aggregation of the beads. Afterwards the immobilized biocatalyst beads were filtered, washed three times with 1 mL deionized water and stored at 4°C prior to use. The CaCl_2_ solution and the filtered water were stored to calculate the immobilization efficiency by measuring the recovered protein. The percentage of immobilization was calculated as follows:

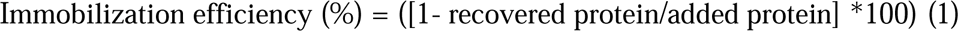

#### 2.4.2. Immobilization of CtNDT on Chitosan Beads

*Ct*NDT was immobilized by direct adsorption and covalent binding into chitosan beads. The beads were crafted by preparing a 3 % high molecular weight chitosan solution in acetic acid (1.5 %). 3 g of chitosan powder were dissolved in 100 mL of acetic acid under magnetic stirring until a homogenous solution was formed. Chitosan beads were formed by extruding the solution through a syringe needle (0.8 x 40 mm) into a coagulant solution comprised of 1 M NaOH in a constant stirring. These freshly formed beads were allowed to rest under magnetic stirring for a period of 48 hours at a temperature of 4 °C. After hardening, chitosan beads were filtered and washed 3-5 times to reach neutral pH, and finally stored at 4°C for further application.

##### 2.4.2.1. Direct adsorption

For the direct adsorption of CtNDT onto chitosan beads, a suspension comprising 250 μg of enzyme per gram of chitosan beads was stirred at 100 rpm at 25 °C for 3 hours. The suspension was meticulously prepared using 7.5 mL of phosphate buffer (5 mM, pH 7.5) per gram of the carrier material. Afterwards, the beads were filtered, washed three times utilizing an equivalent volume of buffer and stored at 4 °C for further use. The calculation of the immobilization percentage was carried out using the same methodology as described above.

##### 2.4.2.2. Covalent binding

CtNDT was immobilized by covalent binding into chitosan beads using glutaraldehyde as a crosslinking agent. Firstly, 0.5 g of chitosan beads were activated by mixing with 5 mL of 5 % glutaraldehyde solution for 1 h at 25 °C in a shaking incubator (100 rpm). Then, surplus glutaraldehyde was removed by rinsing the beads with deionized water. Once the beads were activated, covalent immobilization was achieved by following the same steps as described for direct adsorption immobilization.

#### 2.4.3. Covalent immobilization of CtNDT_Y7F_A9S_ and BpNDT on Chitosan Beads

CtNDT_Y7F_A9S_ and *Bp*NDT were covalently immobilized through a direct binding process, mirroring the procedure employed for *Ct*NDT. The initial step involved the chemical activation of chitosan beads using a 5 % glutaraldehyde solution. Once the beads were washed, two different suspensions were used for the covalent immobilization: 250 μg of enzyme per gram of beads for CtNDT_Y7F_A9S_, and 1000 μg of enzyme per gram of beads for *Bp*NDT. As a final step, each suspension solution was used for the covalent immobilization utilizing a procedure identical to the one employed for immobilizing *Ct*NDT within the chitosan beads.

### 2.5. Characterization of enzyme immobilized on Chitosan beads

#### 2.5.1. N-Deoxyribosyltransferase assays for free and immobilized enzymes

The activity of 2′- deoxyribosyltransferases was assessed through a reaction involving a 1 mM nucleoside substrate and 1 mM nucleobase substrate. To determine the specific activity of *Ct*NDT, 2′-deoxyadenosine production from 2′-deoxyguanosine and adenine was used, for the activity of *Ct*NDT_Y7F_A9S_, adenosine production from guanosine and adenine was used, and for *Bp*NDT activity 2′-deoxythymine from 2′- deoxyadenosine and thymine was employed.

Variations in substrates, alongside temperature (ranging from 25 °C to 55 °C) and pH (30 mM CHES, MES, HEPES, within the range of 6.5 to 8.5), were introduced in alignment with the specific enzyme under study (as detailed in Table S1 and Figure S1). Using stablished conditions for each enzyme of interest, after 15 min of incubation the reaction was stopped by the addition of 1000 μL of cold methanol and heating for 5 min at 100 °C as described for other NDTs^20^. Then, 100 μL of quenched reactions were placed into a 96-well round bottom microplate (Agilent Technologies) and 10 μL of each sample were analysed by High-Performance Liquid Chromatography (HPLC, Shimatzu CBM- 20A) with a HSS T3 column 2.5 mm, 50 mm X 4.6 mm (Waters^TM^) monitoring absorbance at 260 nm. A gradient elution was employed to separate reactants and products using buffer A (H_2_O + 0.1% TFA) and buffer B (ACN + 0.1% TFA), from 0-10 mins, 99% to 85% buffer A and 1% to 15% buffer B, from 10-15 mins, 85% to 0% buffer A and 15% to 100% buffer B. The oven temperature was set at 40 °C and the flow rate was 1 ml/min. The column was equilibrated with buffer A: buffer B (1:99) for at least 15 mins before each injection. Calibration curves were performed by integrating peak areas for compound standards from 0.0 to 2.5 mM. One international unit (IU) is defined as the amount of enzyme needed to convert 1 µmol of nucleoside substrate into product per minute.

#### 2.6.2. FTIR

FTIR-ATR analysis (Shimadzu IR Affinity 1S IR Spectrometer – for solid or liquid samples) was performed to characterize chemical functional groups present in alginate beads and chitosan beads before and after each immobilization process. In total, 32 scans were collected within a range of 4000- 400 cm^-1^ with a 1 cm resolution. As sample treatment, beads were dried using N_2_ to remove unbound water and then an ATR-diamond crystal additament was implemented for direct measurements.

#### 2.5.3. Recyclability of immobilized proteins

Immobilized proteins (1.8-18 μg) were evaluated for nucleoside 2’-deoxyribosyltransferase activity after sequential reaction cycles reutilizing the same immobilized enzyme catalyst. After each cycle, biocatalyst was filtered and washed to eliminate residual substrates bond to the support, while the supernatant was treated as previously described and analyzed by HPLC to quantify nucleoside production by HPLC. The recyclability of the immobilized proteins was determined by comparing the nucleoside conversion in each consecutive reaction cycle to the results obtained in the first cycle.

#### 2.5.4. Storage Stability of immobilized proteins

The immobilized proteins were stored for 30 days at 4 °C and samples were periodically removed to measure activity every 7 days according to standardized protocols. The storage efficiency was defined as the ratio of free or immobilized enzyme activity after storage to their initial activity.

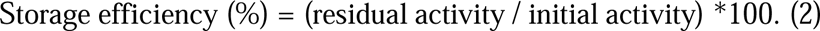

### 2.6. Biosynthesis of 2′-fluoro-3-deoxyadenosine

The synthesis of the nucleoside analogue was performed using a 10 mL reaction mixture consisting of 1 mM of 3′-deoxyadenosine, 1 mM of 2- fluoroadenine, and 5 µM of immobilized enzyme (Chi/CtNDT_Y7F_A9S_Cv). This mixture was suspended in a 30 mM CHES, MES, HEPES buffer (pH 6.5) and heated to 55 °C for 60 minutes. The mixture was filtered to recover the biocatalyst for further reaction cycles, and the flow through was freeze-dried and resuspended using 1 mL of water for subsequent characterization. A total of 7 cycles were performed using the immobilized enzyme.

### 2.7. Characterization of synthetized 2**′**-fluoro-3-deoxyadenosine

#### 2.7.1. HPLC purification

A concentrated solution (10 µL) was injected onto a HSS T3 column 2.5 mm, 250 mm, X 4.6 mm (Waters^TM^) pre-equilibrated with buffer A: buffer B (1:99) for at least 30 mins before each injection (buffer A: 10 mM trimethylammonium acetate, pH 7; buffer B: ACN + 0.1% TFA), while monitoring absorbance at 260 nm. A gradient elution was employed to separate reactants from products using buffer B, from 0-40 mins, 99% to 75% buffer A and 1% to 25% buffer B, from 40- 60 mins, 75% to 0% buffer A and 25% to 100% buffer B. The oven temperature was set at 40 °C and the flow rate was 1 mL/min. Retention times were 3′-deoxyadenosine (19.4 min), 2-fluoroadenine (15.2 min), adenine (12.7 min), and 2′-fluoro-3-deoxyadenosine (22.2 min). Retention time of 2′-fluoro-3-deoxyadenosine was differentiated from the substrates and adenine by comparison to commercial standards. Once retention times were determined, the remaining of the reaction solution was injected and fractions from 21-23 min were collected to isolate 2-fluoro-3′-deoxyadenosine. Fractions were freeze dried, weighed and storage at -20 °C for further analysis.

#### 2.7.2. High resolution mass spectrometry

Mass spectrometry was employed to analyse reaction solutions and isolated 2′-fluoro-3-deoxyadenosine. Samples were injected onto a YMC Triart C18 trap column (12nm, 3uM, 0.3 x 0.5mm) coupled to a YMC Triart C18 analytical column (12nm 3um 0.3 x 150mm) column, using a Eksigent Ekspert nanoLC 425. Compounds were eluted with a gradient between solvents A (water + 3% acetonitrile with 0.1 % formic acid) and B (acetonitrile with 0.1 % formic acid) as follows: 3-95% acetonitrile in 6 min followed by 95% acetonitrile for 2 min, and column re-equilibration with solvent A. The eluate was sprayed into a TripleTOF 6600 electrospray mass spectrometer (ABSciex, Foster City, CA) acquiring MS for 250 msec of accumulation time from m/z 120-1000.

#### 2.7.3. Fluorine nuclear mass spectrometry characterization

Fluorine NMR (^19^F NMR) was applied for the study of the local chemical environment and structure of the biosynthesized molecule. The isolated 2′-fluoro-3-deoxyadenosine as well as an amount of 2-fluoroadenine were dissolved in deuterated dimethyl sulfoxide (DMSO-d6) for its analysis in a Bruker AVII 400 MHz spectrometer equipped with a BBFO probe. 19F NMR was obtained at 376.5 MHz. The experiment was conducted at 298K with a relation delay of 1.5s and 64 scans. Data were processed using MestReNova 15.0.0- 34764.

## 3. RESULTS AND DISCUSSION

### 3.1. Immobilization of *Ct*NDT

Utilizing NDTs while free in solution limits their application due to high cost and complex processing to produce, recover and recycle enzyme^35^. To overcome these issues, different immobilization techniques were explored to prepare nanobiocatalysts of the multimeric NDT from *Chroococcidiopsis thermalis* (CtNDT), including encapsulation in alginate beads (AlgB), physical adsorption, and covalent attachment onto chitosan (Chi) (Figure 1).

**Figure 1.**
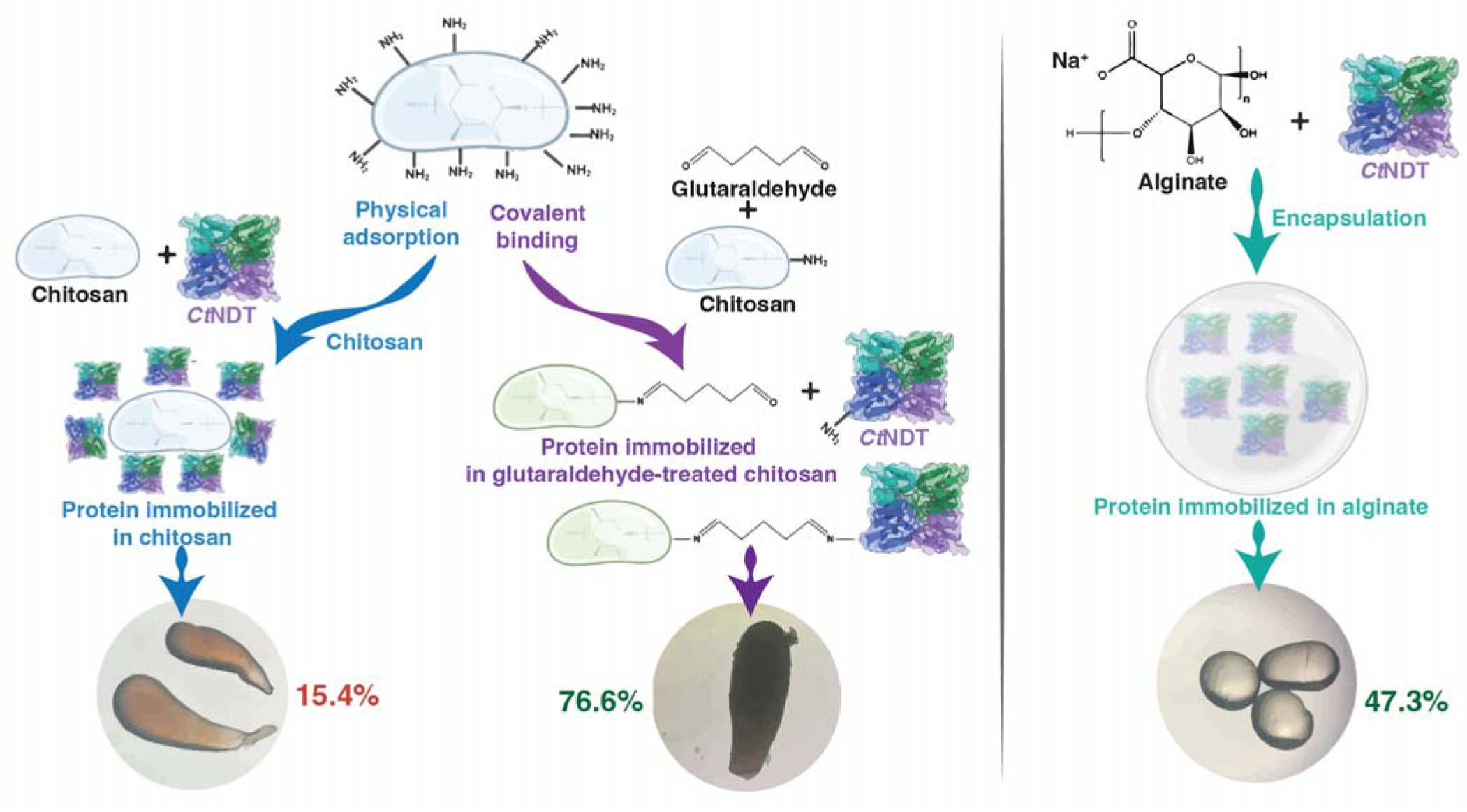
Schematic representation of the strategies employed to immobilize CtNDT. Left: Using chitosan for physical or covalent attachment after glutaraldehyde treatment, immobilization fraction was 15.4% and 76.6% respectively; right: encapsulation into alginate beads, and immobilization fraction was 47.3% . Images in the bottom were obtained using a standard microscope with total magnification of 100x.

#### 3.1.1. Encapsulation of CtNDT in Alginate Beads (AlgB)

*Ct*NDT was immobilized onto AlgB by the direct encapsulation of the enzyme within the polymeric structure of alginate. Encapsulation immobilization consists in the confinement/entrapment of the enzyme inside a spherical semi-permeable membrane by using a two- step process, established by mixing the enzyme with a polymeric monomer, followed by a polymerization reaction^36^. Since *Ct*NDT is stable at pH 7.5 and 25 °C^37^Click or tap here to enter text., the immobilization procedure was carried out with these reaction conditions, monitoring CtNDT’s specific activity by the production of 2′-deoxyadenosine from 2′-deoxyguanosine and adenine substrates. As shown in Table 1, almost half of the CtNDT employed was trapped inside AlgB, but a significant portion of the immobilized enzyme lacked activity. It is important to point out that prior to this work no NDTs were successfully immobilized through encapsulation within a polymeric matrix. Recently, Rivero et al., reported the entrapment of the NDT enzyme from *Lactobacillus delbrueckii* using biomimetic silica nanoparticles (SiBio), however, a significant loss of catalytic activity and immobilization percentage was observed in this system^38^.

**Table 1.**
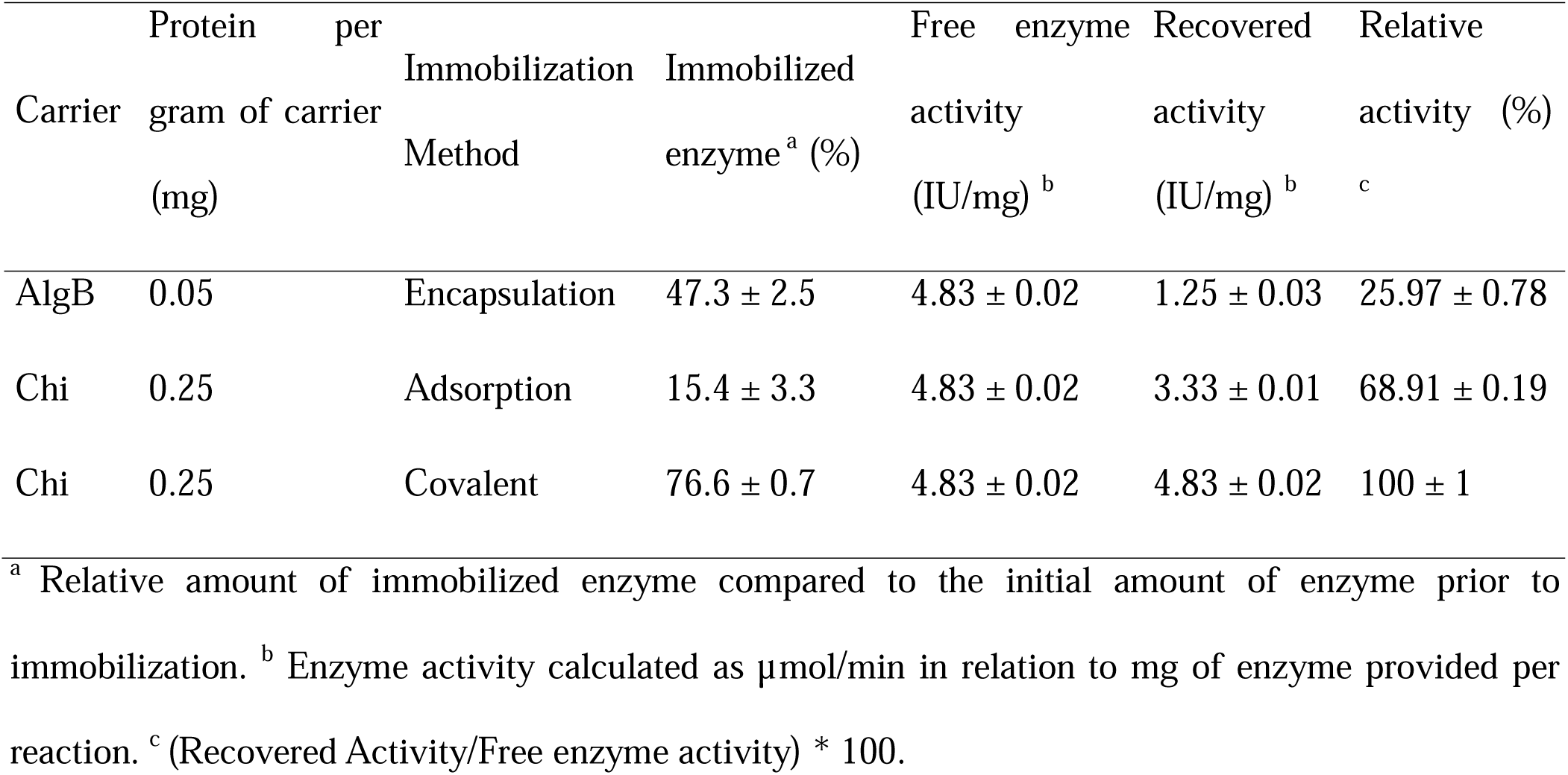
Immobilization of CtNDT on Alginate Beads and Chitosan by different immobilization procedures.

High immobilization percentages accompanied by loss of activity have been reported for the encapsulation of enzymes within alginate beads or similar polymeric matrixes, including the immobilization of glycoenzymes, tyrosinases and glucosidases in sodium alginate beads^39,40^. The low enzymatic immobilization percentage and lower activity of the CtNDT onto AlgB may be attributed to the heterogenous gel structure, enzyme leakage and poorly substrate diffusion through the microporous structure of alginate beads^41^. Despite lower activity compared to its free form, AlgB immobilized CtNDT was tested for recyclability to enable a comparison with non-immobilized enzyme.

#### 3.1.2. Immobilization of CtNDT into Chitosan

We evaluated how CtNDT performed when immobilized by physical adsorption and covalent attachment into chitosan. Physical adsorption consists of the non-covalent interaction between enzyme and carrier, mediated by the adsorption of the enzyme into the matrix surface through ionic interactions, hydrogen bonding, and Van der Waals forces ^36^. Covalent immobilization of proteins occurs via the multipoint covalent binding between functional groups of the carrier and amino groups from the protein^42^.

The physical adsorption of CtNDT onto chitosan beads occurred with low immobilization efficiency (15.4 %, Table 1), and therefore most of the protein could not adsorb to the surface of chitosan. In terms of activity, adsorbed protein retained 68.91 % of specific protein activity compared to its free form. Different NDT species have been immobilized by ion exchange mechanisms, including the immobilization of *Lactobacillus animalis* NDT into IDA-Agarose, boronate Agarose, DEAE-sepharose, and Q-agarose ^43^, in which immobilization percentage was greater than the one obtained here (41- 93 % immobilization), however, in that study much smaller amounts of enzyme were employed in the adsorption process, which might suggest that our low immobilization yield was due to saturation of chitosan surface. Furthermore, the reported activity was very low or undetectable in comparison to the activity observed after the immobilization of CtNDT over chitosan (0.027 IU/g compared to 0.12 IU/g). No loss of catalytic activity was reported by the immobilization of NDT from *Bacillus psychrosaccharolyticus* (BpNDT) onto polyethyleneimine-agarose support (PEI-agarose)^20^. However, this preparation suffered from low stability and subsequent loss of activity due to enzyme leakage, suggesting other immobilization techniques such as covalent immobilization could overcome this inherent disadvantage of physical adsorption.^44^

Aiming to improve support attachment, we performed the covalent immobilization of *Ct*NDT using glutaraldehyde activated chitosan. The immobilization process achieved 76 % of immobilization efficiency and led to negligible effects on enzyme activity. Immobilization by covalent binding onto chitosan might take place by the activation of chitosan with glutaraldehyde through amine groups in chitosan, followed by the formation of a Schiff’s base between activated carrier and proteins (Figure 1)^42,45^. Due to the relatively high immobilization yield compared to other NDTs, and its high enzymatic activity, covalent immobilization onto chitosan was also applied for the immobilization of different enzymes, including a novel CtNDT mutant and BpNDT.

Although covalent immobilization has been previously reported for the NDT enzymes from *Lactobacillus delbrueckii*^23^*, Trypanosoma brucei*^46^*, and Lactobacillus animalis*^47^, reducing enzyme leakage^36^, these attempts also resulted in significant loss of activity. NDT enzymes immobilized into nanomaterials reported losses of 66 % and 50 % of protein activity after covalent immobilization^42,48^. On the other hand, the preparation of a robust biocatalyst by covalent immobilization into silica biomimetic nanoparticles^23^ using a two-step process (glutaraldehyde activation followed by covalent binding) with the NDT from *Lactobacillus delbrueckii*, maintained enzymatic activity but suffered from low immobilization percentage.^38,27,30^

#### 3.1.3. Covalent immobilization of CtNDT_Y7F_A9S_ and BpNDT onto chitosan

Since covalent immobilization was determined as the preferable option for CtNDT, the same methodology was applied for the attachment of CtNDT_Y7F_A9S_, a double mutant of CtNDT with improved catalytic turnover when utilizing ribonucleoside substrates.

Additionally, BpNDT, which was previously immobilized^49^ and shown to suffer from support leakage^20^ was immobilized onto chitosan beads to test whether leakage could be overcome by this strategy. As anticipated, CtNDT_Y7F_A9S_ (Table 2) showed similar immobilization efficiency (between 70-80 %) as the wild-type enzyme. Specific activity was decreased in the chitosan beads in comparison to its free form and to wild type protein (79.6 % activity, while no loss occurred in the wild type). Despite this decrease in activity relative to wild-type CtNDT, the immobilization of CtNDT_Y7F_A9S_ is comparable to results obtained with other biocatalysts reported^20,48^. In contrast, BpNDT had high immobilization percentage but only 26 % of the immobilized enzyme remained active for the production of 2′- deoxythymine. This is in line with what was described for the immobilization of BpNDT over PEI-functionalized carriers, where complete immobilization of the enzyme was achieved with 56 % of activity retention^20^. However, a multistep immobilization process was applied, requiring adsorption, crosslinking, and chemical reduction to achieve covalent immobilization, ultimately leading to approximately 40 % loss of catalytic activity after 24 h.

**Table 2.** Covalent immobilization of *CtNDT_Y7F_A9S_* and *Bp*NDT onto chitosan beads.

| Carrier | Protein per gram of carrier (mg) | Enzyme | Immobilized enzyme <sup>a</sup> (%) | Free enzyme activity (IU/mg) <sup>b</sup> | Recovered activity (IU/mg) <sup>b</sup> | Relative activity (%) <sup>c</sup> |
| --- | --- | --- | --- | --- | --- | --- |
| Chi | 0.25 | CtNDT<sub>Y7F_A9S</sub> | 72.3 ± 2.1 | 0.16 ± 0.08 | 0.13 ± 0.01 | 79.6 ± 0.4 |
| Chi | 1.00 | BpNDT | 95.0 ± 2.3 | 0.38 ± 0.03 | 0.099 ± 0.008 | 26.1 ± 2.5 |

<sup>a</sup> Relative amount of immobilized enzyme in comparison to enzyme prior to the immobilization process. <sup>b</sup> Enzymatic activity calculated as $\mu\text{mol}/\text{min}$ in relation to mg of enzyme provided per reaction. <sup>c</sup> (Recovered Activity/Free enzyme activity) \* 100.

### 3.2. Physicochemical characterization of biocatalysts by FTIR

FTIR characterization was performed to identify functional groups present in both non-modified immobilization matrixes and final biocatalysts. The characteristic FTIR spectrum of alginate beads (AlgB) is shown in Figure 2A. The polymeric matrix is composed of a broad band centered at 3333 cm^-1^, corresponding to hydroxyl (-OH) stretching; a low intensity band at 2920 cm^-1^ from -CH_2_ stretching, medium and low intensity peaks at 1640 cm^-1^ and 1418 cm^-1^, which are attributed to carboxylic (-COO) asymmetric and symmetric stretching modes, respectively, and low bands between 1025 cm^-1^ and 1090 cm^-1^ attributed to C-O-C stretching ^50^ After the addition of *Ct*NDT onto AlgB, a significant difference is observed between spectra. Some signals are present in both spectrum, including the 3333 cm^-1^, and 1640 cm^-1^, however, these peaks might be overlapped by -OH and -NH stretching (3333 cm^-1^); and - COO stretching and amine N-H bending (1640 cm^-1^). This is supported by the presence of two medium bands centered between 950 cm^-1^ and 1025 cm^-1^, corresponding to C-N stretching ^51^. Moreover, the low intensity band located at 820 might be related to N-H wagging vibrations ^52^. In this context, the presence of different signals attributed to functional groups presented in enzymes, such as amide I, amide II and amines confirm the immobilization of *Ct*NDT onto the AlgB matrix. *Ct*NDT was immobilized into chitosan beads by two different methodologies, physical adsorption, and covalent binding. Figure 2B shows the characteristic FTIR spectrum of chitosan beads (black spectrum). A broad band located at 3300 cm^-1^, which might correspond to O-H stretching vibrations of hydroxyl groups presented om the saccharide units. In addition, lower signals were identified, corresponding to C=O stretching of amide I group (1645 cm^-1^), and lower bands between 1050 – 1090 cm^-1^ corresponding to C-O-C stretching vibrations of the glycosidic linkages ^51,53^). Furthermore, the presence of the different proteins in chitosan beads, including CtNDT, CtNDT_Y7F_A9S_, and BpNDT were identified by characteristic peaks. Signals between 1645 cm^-1^ and 1660 cm^-1^ are attributed to C=O stretching from Amide I, however, since chitosan also contains this group only a slight shift can be observed, which can be attributed to the low enzyme loaded in comparison to the chitosan of high molecular grade utilized. The same signals between 1050 cm^-1^ and 1090 cm^-1^ corresponding to C-O-C stretching from chitosan matrix are observed, which confirms that the chitosan structure is still present after protein modification [3]. Finally, a low signal at 2980 can be observed for those biocatalysts that suffered covalent immobilization, this peak is attributed to C-H stretching vibrations of -CH3 and -CH2. In this context, this signal could be indicative of the aliphatic side chains of amino acids in the protein, which suggests that a protein modification could be introduced or affected by these -CH_3_ and -CH_2_ ^51,54^.

**Figure 2.**
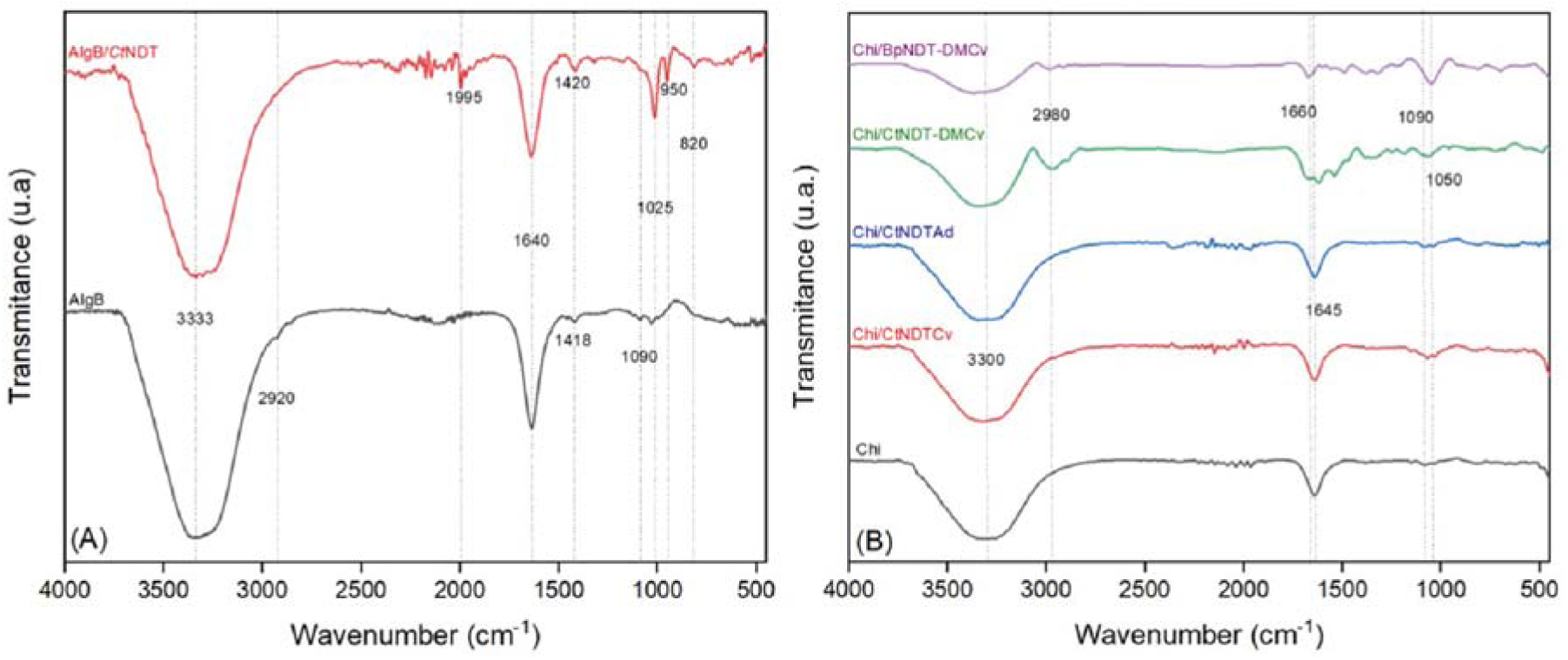
(A) FTIR spectra of alginate beads (AlgB) and encapsulated *Ct*NDT in alginate beads (AlgB/CtNDT); (B) FTIR spectra of chitosan beads (Chi), covalently immobilized CtNDT onto chitosan (Chi/CtNDTCv), adsorbed CtNDT onto chitosan (Chi/C*t*NDTAd), covalently immobilized BpNDT onto chitosan beads (Chi/BpNDTCv).

### 3.3. Recyclability of immobilized enzymes

Each biocatalyst was tested across multiple cycles to assess their recyclability during nucleoside production. These catalysts included entrapped CtNDT onto AlgB, adsorbed CtNDT onto the chitosan, and CtNDT, CtNDT_Y7F_A9S_, and BpNDT covalent attached onto chitosan beads. Each recycling process was halted either when the material showed evident damage, complete enzymatic deactivation, or reached a maximum of 20 cycles as a limit.

As aforementioned, soluble enzymes pose challenges for recycling which complicates their use in industrial applications.^40^ This issue can be addressed by enzyme immobilization. Although immobilization often leads to partial or significant loss of enzyme activity, the ability to recycle the enzymes can greatly mitigate this drawback.^44^ As shown in Figure 3A, reusing each biocatalyst overcame this limitation as only a few cycles (ranging from 1 to 5, depending on the enzyme) were required to achieve the same nucleoside production in comparison to enzymes free in solution. We were able to recycle the biocatalysts Chi/*Bp*NDTCv, AlgB/*Ct*NDT, and Chi/*Ct*NDTAd for 7, 11, and 12 consecutive cycles, respectively. This recycling led to an increase in the relative nucleoside production of 140 %, 212 %, and 376 %, respectively. More pronounced improvement was observed when the NDT biocatalysts Chi/*Ct*NDTCv and Chi/ CtNDT_Y7F_A9S_ Cv were recycled. Both enzymes were successfully applied for 20 consecutive cycles, resulting in a relative nucleoside production increase of more than 1000% in comparison to the same enzymes free in solution.

**Figure 3.**
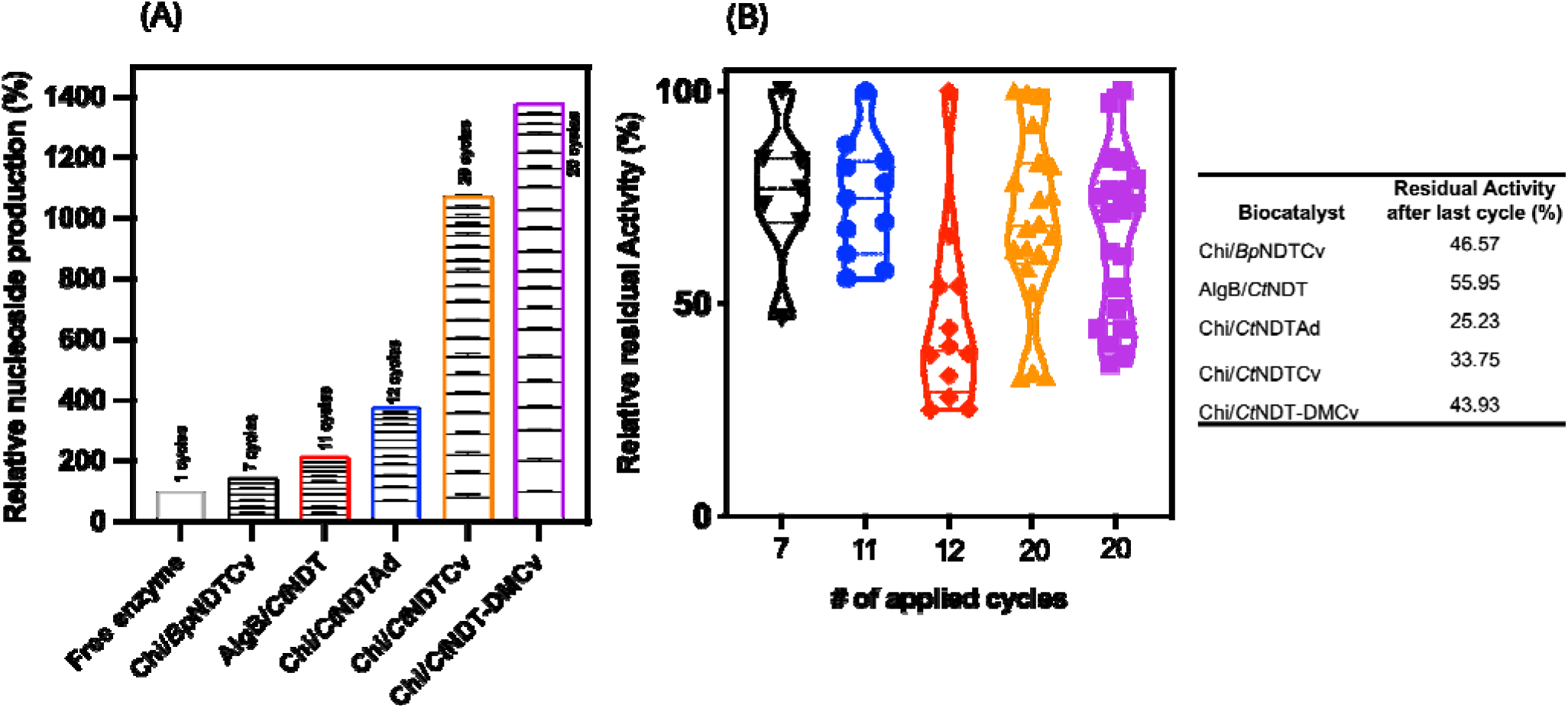
Relative nucleoside production ^a^ (A), in which blocks are consecutive cycles; and relative residual activity (B) ^b^ represented by violin plots showing the distribution of residual enzyme activity density, in which each dot represents the mean enzyme activity for each cycle. (^a^Relative to the production of NAs by free enzyme system. ^b^Relative to initial biocatalysts activity). Abbreviations and colors: BpNDT covalently immobilized into chitosan beads (Chi/BpNDTCv, black), CtNDT entrapped into alginate beads (AlgB/CtNDT, red), CtNDT adsorbed onto chitosan beads (Chi/CtNDTAd, blue), CtNDT covalently immobilized into chitosan beads (Chi/CtNDTCv, orange), and CtNDT_Y7F_A9S_ covalently immobilized into chitosan beads (Chi/CtNDT-DMCv, purple).

In addition, Figure 3B depicts the maintenance of enzymatic activity during the recycling process. We observe the percentage of remaining activity as well as the distribution of cycles throughout the process, with 100% set as the activity observed during the first cycle. The adsorption immobilization of *Ct*NDT onto chitosan beads resulted in the biocatalyst with the highest loss of enzymatic activity (approximately 75%). Furthermore, as noted, at the beginning of the recycling cycle the activity decreased abruptly, which could be attributed to enzyme leakage from the material, as has been reported for other NDT immobilization methods by adsorption^20,43^. In contrast, encapsulation immobilization was a better option in terms of retaining enzymatic activity. This agrees with the two main characteristics previously reported for encapsulation: 1) structural changes are not occurring in the enzymes and 2) the polymeric matrix avoids enzymatic leakage.^36^ The remaining activity for biocatalysts made by covalent immobilization was between 33.75 % and 46.57 %. In these cases, most activity deactivation is likely due to structural modifications made by the addition of covalent bonds

### 3.4. Storage stability of immobilized enzymes (**Figure 4**)

The storage stability at 4°C was tested for all prepared biocatalysts and compared with their respective free enzyme forms. Figure 4A shows the residual activity (%) of the immobilized enzymes for samples measured every 7 days within a period of 28 days. As could be expected, the activity of the NDT enzymes decreases over time when stored at 4°C instead of -80 °C, but significant activity was retained for all catalysts. Figure 4A shows the behavior obtained for the immobilized enzymes, in which encapsulation and covalent immobilization of CtNDT and CtNDT_Y7F_A9S_, respectively, presented higher stability retaining ∼75 % of their initial activity after 28 days. Comparing to their free form (Figure 4B), CtNDT immobilization exhibited similar results when immobilized by encapsulation, but whowed significant loss of activity when adsorption was applied (about 75 %). Activity of CtNDT_Y7F_A9S_ was improved by immobilization, showing higher activity than its free form after 28 days of storage at 4°C. *Bp*NDT immobilization led to small losses of activity compared to its free form (<10 %). Different results have been published for the storage stability of immobilized NDT enzymes. Stability of *Ld*NDT immobilized by adsorption onto PEI supports had complete activity loss after 2 h in DMF solution^20^. Moreover, as observed here, others reported that encapsulation was the better enzyme immobilization method in terms of storage stability, the entrapment of *Ld*NDT onto SiGPEI supports resulted in stable residual activity after 30 days at 4°C.^38^

**Figure 4.**
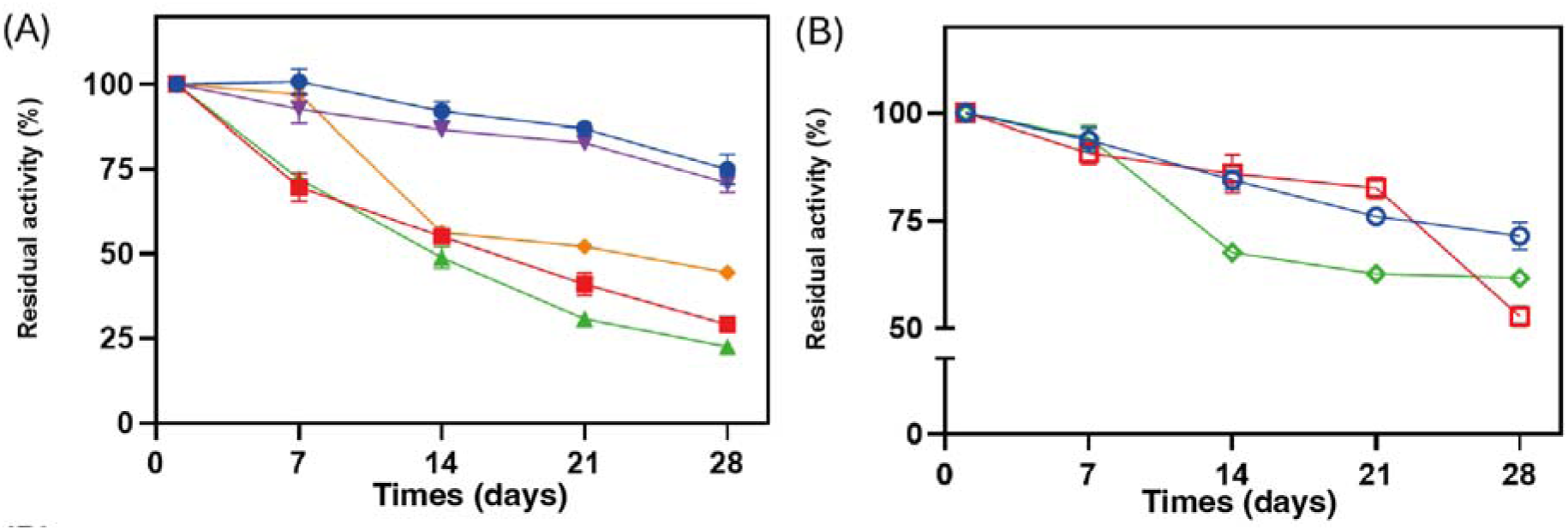
Storage stability at 4 °C. (A) immobilized biocatalysts, AlgB/CtNDT (●), Chi/CtNDTCv (▪), Chi/CtNDTAd (▴), Chi/BpNDT (♦), and Chi/ CtNDT_Y7F_A9S_ Cv (▾). (B) free enzymes, CtNDT (○), CtNDT_Y7F_A9S_ (□), and BpNDT (◊). with the support, differing from enzyme leakage encountered in adsorption strategies.^54^

### 3.5. Biosynthesis and characterization of 2-fluoro-3′-deoxyadenosine

The biosynthesis of the nucleoside analogue 2-fluoro-3′-deoxyadenosine (Scheme 1) was performed during 7 consecutive cycles by CtNDT_Y7F_A9S_ covalently immobilized into chitosan beads. 2-fluoropurine nucleosides have been prepared as they were shown to possess activity against human tumor cells.^55^ HPLC was employed for purification, and retention time for 2-fluoro-3′-adenosine was identified as 22 minutes (Fig S2). A total of 5.90 ± 0.46 mg were isolated after purification, corresponding to the production of 219.3 ± 1.9 % compared to the production expected when free enzyme is used a single time (2.7 mg). This corresponds to 5.8 ± 0.1 mg of 2-fluoro-3′-deoxyadenosine per mg of immobilized enzyme.

The isolated product, as well as the reaction mixture were analyzed by high resolution mass spectrometry (Fig S3, expected mass-to-charge ratio (m/z) 270.0997, observed m/z = 270.1006, deviation of 3.33 ppm), confirming the successful biosynthesis of 2-fluoro-3′-deoxyadenosine. The isolated product contained an impurity corresponding to cordycepin (m/z = 252.24 [M + H]). ^19^F NMR confirmed the biosynthesis of 2-fluoro-3′-deoxyadenosine. The spectrum obtained for 2-fluoroadenine (Fig S4) exhibited a signal corresponding to the unique F atom with a chemical shift of -53 ppm. The ^19^F NMR spectrum for the biosynthesized product (Fig S5) exhibit one main peak at -73 ppm and the presence of a small signal at -53 ppm, suggesting the presence of 2-fluoroadenine as a minor impurity. Different chemical shift between signals may be attributed to shielding effects caused by the dissimilarities in molecular structures and local environments. In the case of 2-fluoroadenine, the fluorine atom is in the purine ring, which contains electronegative atoms that promote deshielding, making the fluorine nucleus more exposed to the applied magnetic field and with higher chemical shift. 2-fluoro-3′-deoxyadenosine is a more complex molecule, and the fluorine atom can suffer shielding effects due to electron density from surrounding atoms, compounded by steric effects changing the chemical shift to lower ppm values.^56^ Overall, the shielding effects in 2-fluoro-3′-deoxyadenosine are likely responsible for the observed downfield shift compared to the more deshielded 2-fluoroadenine in a ^19^F NMR spectrum.

**Scheme 1.**
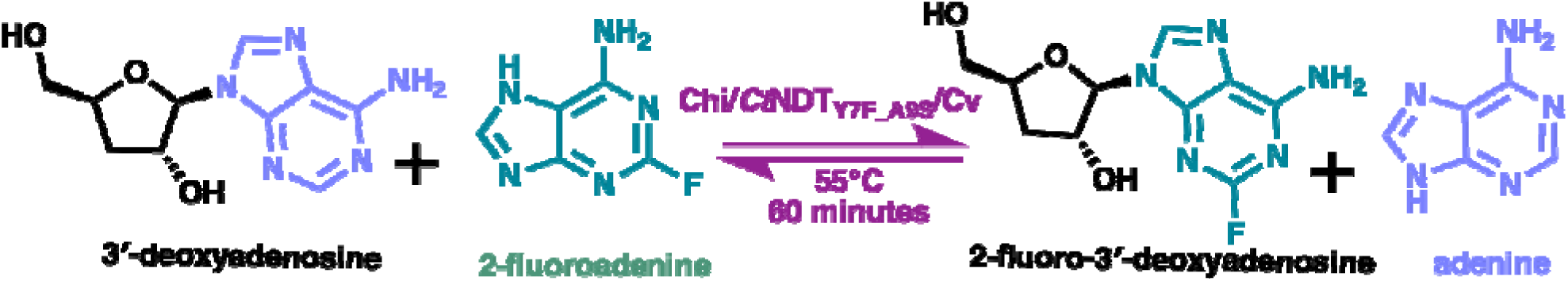
Schematic representation for the biosynthesis of 2-fluoro-3′-deoxyadenosine catalyzed by immobilized CtNDT_Y7F_A9S_.

## 4. CONCLUSIONS

We immobilized NDTs enzymes from *Bacillus psychrosaccharolyticus* (*Bp*NDT) and *Chroococcidiopsis thermalis* (*Ct*NDT) including an improved mutant for the production of ribonucleosides (CtNDT_Y7F_A9S_) onto alginate beads and chitosan using different immobilization methods. Our work provides a detailed method for the utilization and recycling of NDTs for the production of nucleosides with therapeutic potential. All three enzymes immobilized could be recy led in subsequent reaction cycles, and we reported an increment in nucleoside production up to 1000 % when CtNDT_Y7F_A9S_ is covalently immobilized in chitosan. Finally, the immobilized CtNDT_Y7F_A9S_ was used to produce the pharmacologically relevant nucleoside 2-fluoro-3′-deoxyadenosine, with activity as an anti-trypanossomal compound.^33^ We developed a successful strategy to immobilize NDT enzymes, including thermostable variants, with little to no loss in activity after immobilization.

## ASSOCIATED CONTENT

### Supporting Information

The Supporting Information is available free of charge. Supporting Figures and Tables with additional experimental details are available.

## AUTHOR INFORMATION

### Corresponding Author

\***Clarissa Melo Czekster**- *School of Biology, University of St Andrews, Fife KY169ST, U.K;*

### Authors

**Saúl Antonio Hernández Martínez-** *School of Engineering and Sciences, Tecnologico de Monterrey, Monterrey 64849, Mexico*.

**Peijun Tang-** *School of Biology, University of St Andrews, Fife KY169ST, U.K*.

**Roberto Parra-Saldívar-** *Autonomous University of Nuevo Léon, School of Agronomy*.

**Elda M. Melchor- Martínez-** School of Engineering and Sciences, Tecnologico de Monterrey, Monterrey 64849, Mexico.

### Author Contributions

P.T. and S.A.H.M. conducted all experiments. The manuscript was written through contributions of all authors. All authors have given approval to the final version of the manuscript.

## ACKNOWLEDGMENTS

We thank Dr Lynette Alvarado-Ramirez for helpful discussions and suggestions on enzyme immobilization and the Liquid-state NMR facility at the University of St Andrews.

## ABBREVIATIONS

NDT enzymes from *Chroococcidiopsis thermalis* (*Ct*NDT), and *Bacillus psychrosaccharolyticus* (*Bp*NDT), 2 - deoxyadenosine(2 - dAdo), 2 - deoxyguanosine(2 - dGuo), Adenine (Ade), Guanine (Gua), Adenosine (Ado), Guanosine (Guo), liquid-chromatography-mass spectrometry (LC-MS), High performance Liquid Chromatography (HPLC).

## TOC graphic

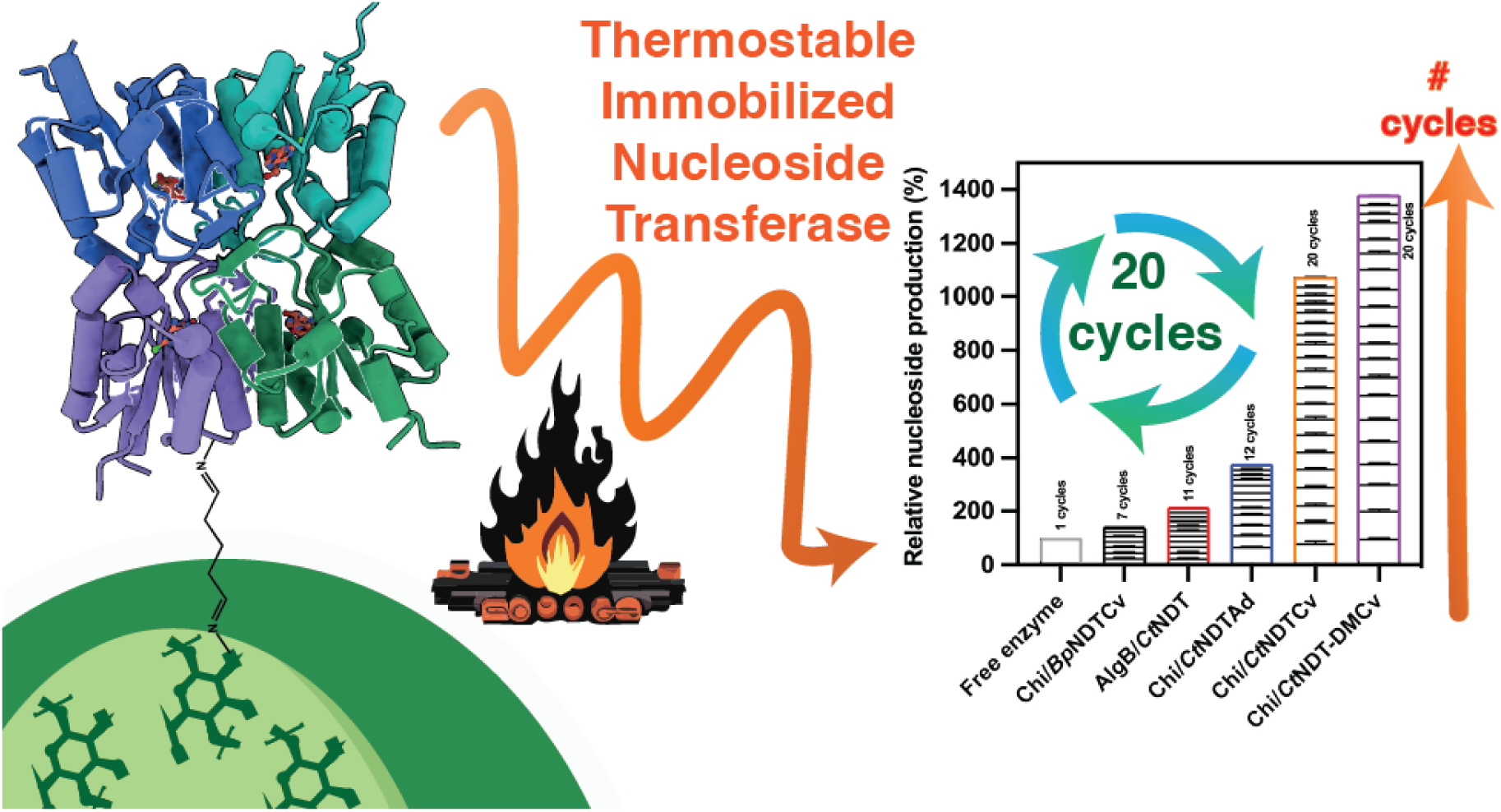

