## Supporting information for "Immobilized Nucleoside 2′-Deoxyribosyltransferases from Extremophiles for Nucleoside Biocatalysis"

**CONTENTS**

**Table S1.** NDTs application for the synthesis of non-natural nucleosides, reaction conditions and HPLC retention times…………………………………………………………………………….2

**Figure S1.** Structures for the enzyme activity assays: 1) 2'- Deoxyguanosine, 2) Guanosine, 3) 2'-Deoxyadenosine, 4) Adenosine, 5) 2'- Deoxythymidine, 6) Guanine, 7) Adenine, and 8) Thymine……….3

**Figure S2.** HPLC chromatogram for the biosynthesis of 2-fluoro-3'-deoxyadenosine catalyzed by immobilized CtNDT-DM……………………………………………………………………………...4

**Table S1.** NDTs application for the synthesis of non-natural nucleosides, reaction conditions and HPLC retention times.

| Enzyme | Reaction | Conditions | Compound | Retention time (min) |
|---|---|---|---|---|
| *Ct*NDT (1.8 µg) | 2'- dGuo + Ade = Gua + 2'- dAdo | 15 min, pH 8.5, 50° C, 180 rpm, 1 mL reaction. | 2'- dGuo | 3.8 |
| | | | Ade | 2.4 |
| | | | Gua | 1.4 |
| | | | 2'- dAdo | 8.2 |
| *Ct*NDT Y7F A9S (18 µg) | RGua + Ade = Gua + RAdo | 15 min, pH 6.5, 55° C, 180 rpm, 1 mL reaction. | RGuo | 3.2 |
| | | | Ade | 2.4 |
| | | | Gua | 1.4 |
| | | | RAdo | 8.0 |
| *Bp*NDT (18 µg) | 2'- dAdo + Thy = Ade + 2'- dThd | 15 min, pH 8.0 25° C, 180 rpm, 1 mL reaction. | 2'- dAdo | 8.7 |
| | | | Thy | 2.2 |
| | | | Ade | 2.6 |
| | | | 2'- dThd | 5.9 |

2'- dGuo (2'- Deoxyguanosine), Ade (Adenine), Gua (Guanine), 2'- dAdo (2'- Deoxyadenosine), RGua (Guanosine), RAdo (Adenosine), Thy (Thymine), and 2'- dThd (2'- Deoxythymidine). For structures FS1

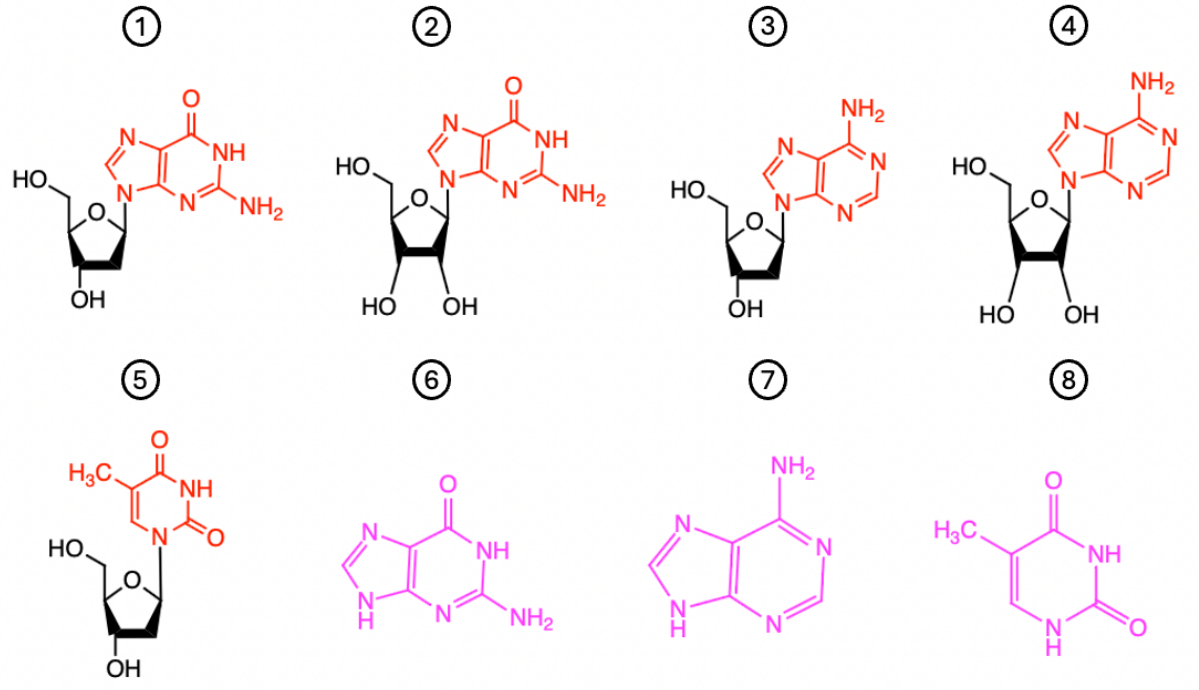

Figure S1. Structures for the enzyme activity assays: 1) 2’- Deoxyguanosine, 2) Guanosine, 3) 2’- Deoxyadenosine, 4) Adenosine, 5) 2’- Deoxythymidine, 6) Guanine, 7) Adenine, and 8) Thymine.

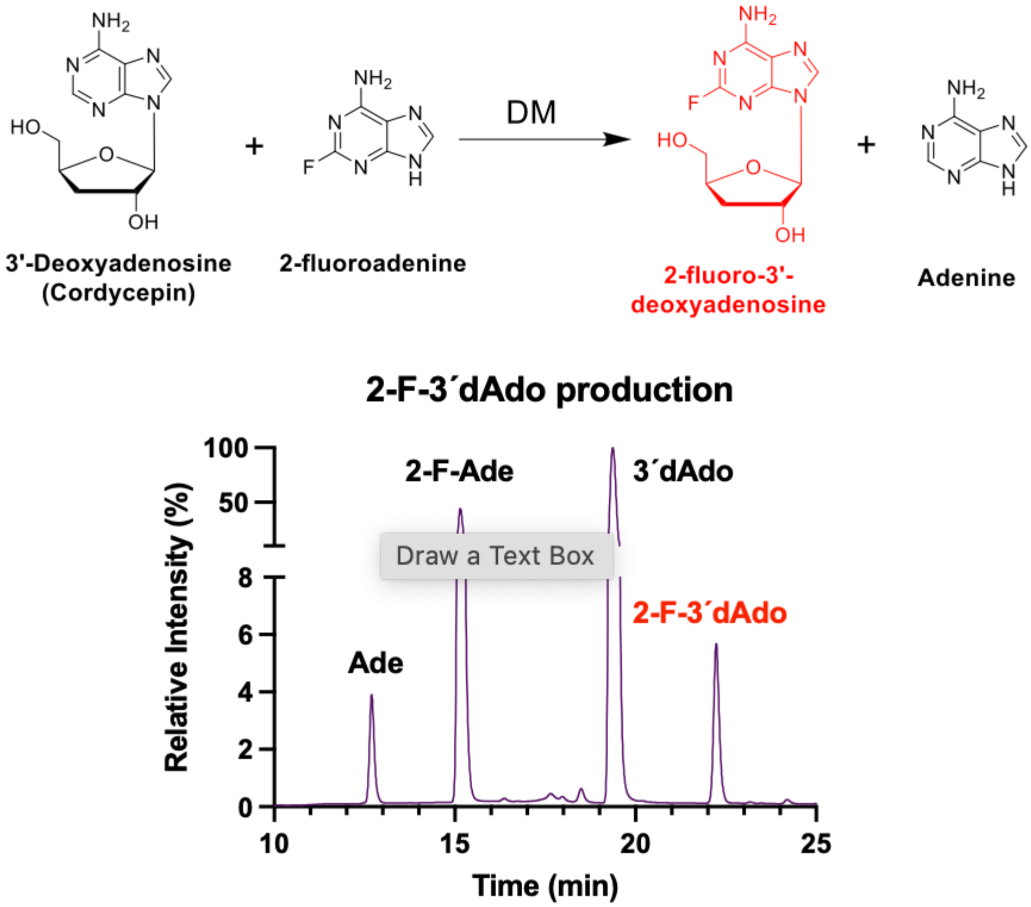

**Figure S2.** HPLC chromatogram for the biosynthesis of 2-fluoro-3'-deoxyadenosine catalyzed by immobilized CtNDT-DM.

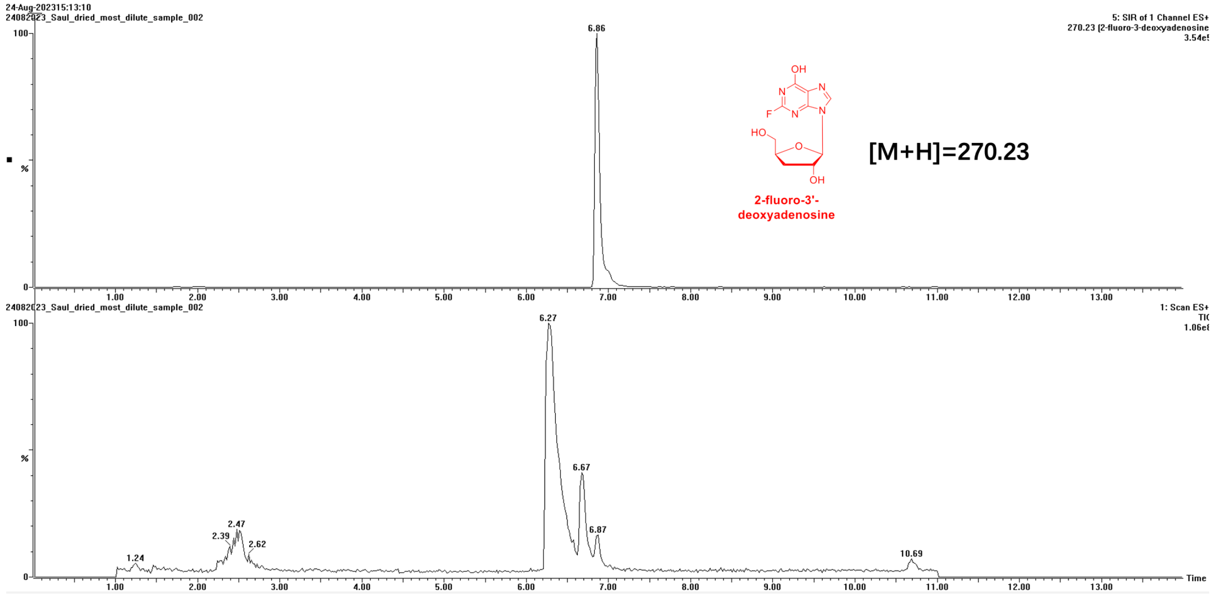

**Figure S3.** Mass Spectrometry for the biosynthesis of 2-fluoro-3'-deoxyadenosine catalyzed by immobilized CtNDT-DM. 3'-deoxyadenosine (19.4 min), 2-fluoroadenine (15.2 min), adenine (12.7 min), and 2'-fluoro-3-deoxyadenosine (22.2 min).

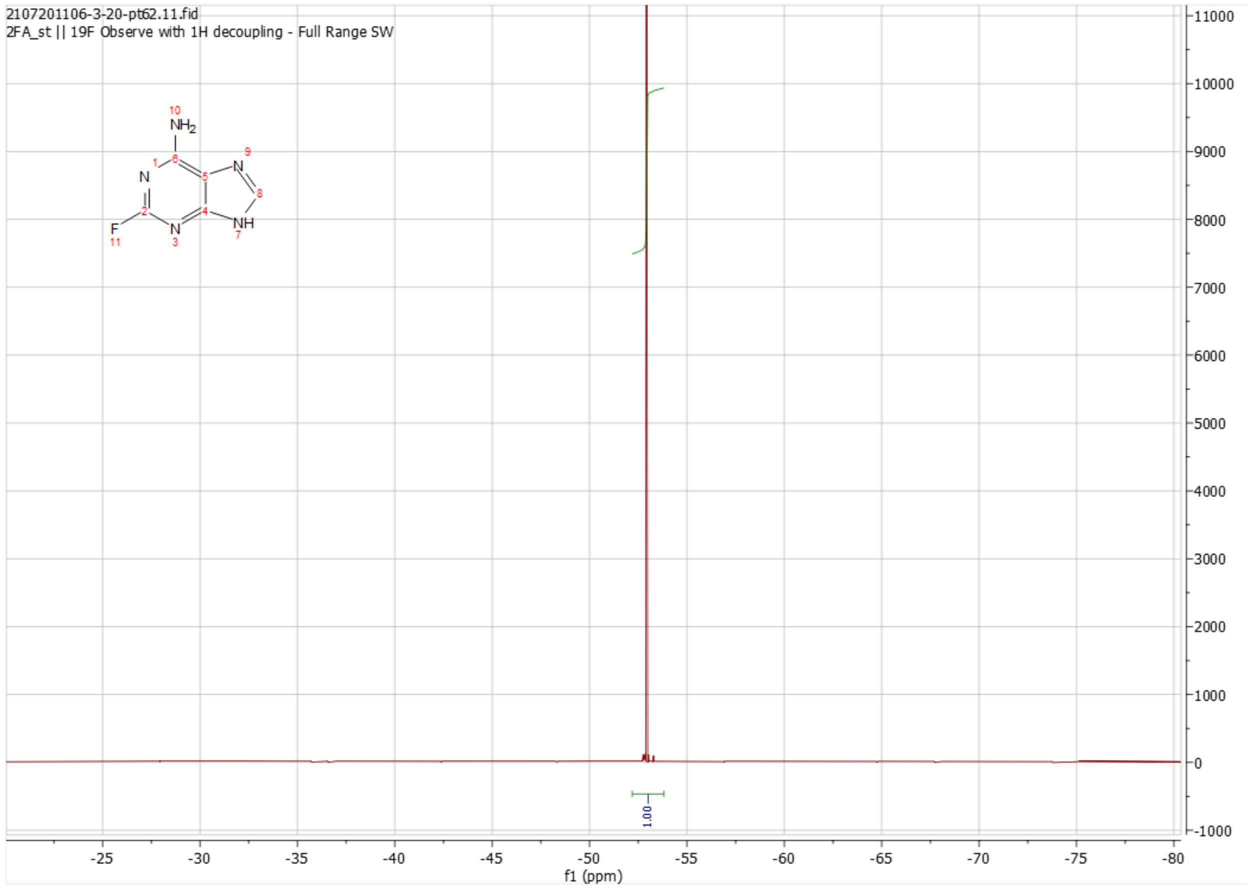

**Figure S4.** 19F NMR spectrum for 2-fluoroadenine standard.

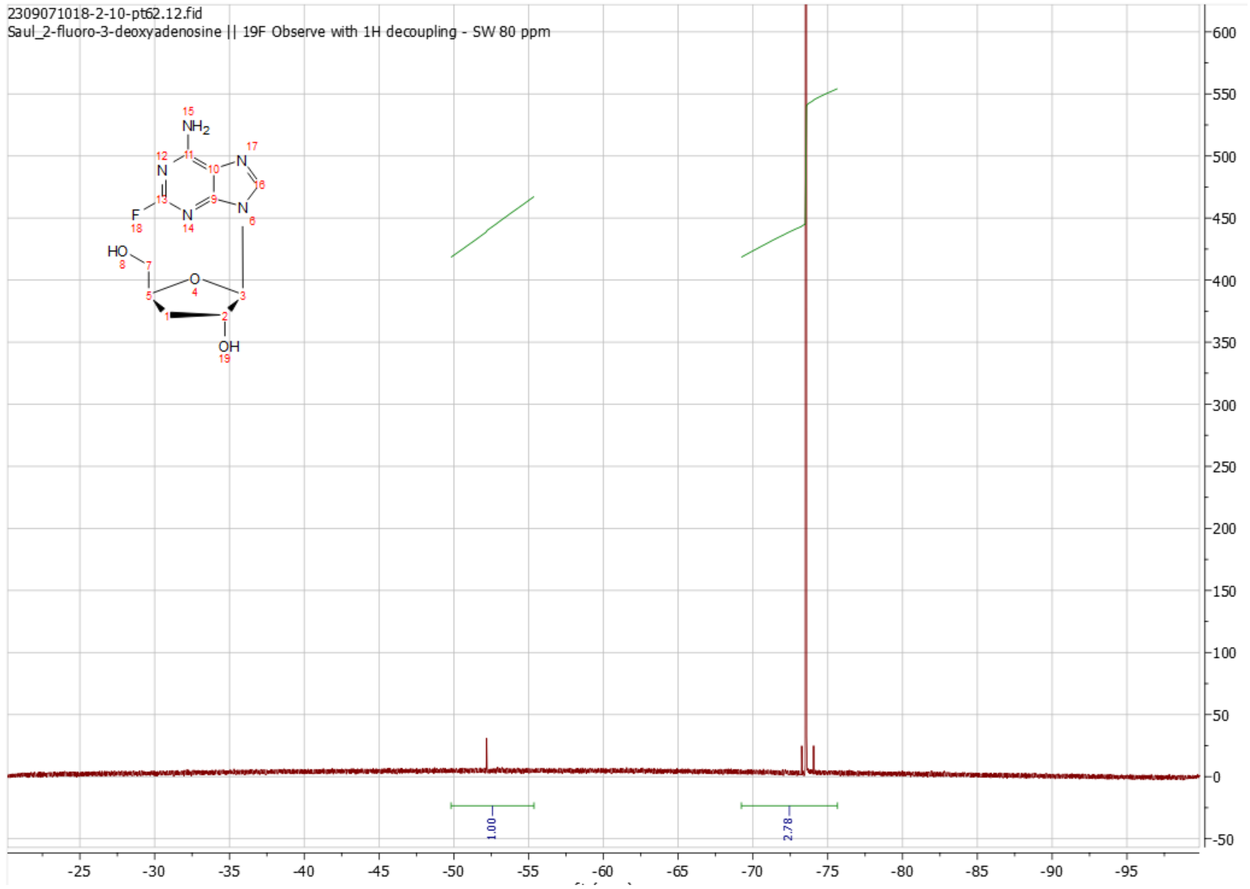

**Figure S5.** $^{19}$F NMR spectrum for the biosynthesis of 2-fluoro-3'-deoxyadenosine catalyzed by immobilized *Ct*NDT-DM.
